# The RNA and protein landscapes of mouse brain organoids

**DOI:** 10.64898/2026.03.13.711293

**Authors:** Anne-Charlotte Fromaget, Céline Gonthier-Guéret, Khadija El Koulali, Sophie Périllous, Nuttamonpat Gumpangseth, Bertille Montibus, Philippe Arnaud, Serge Urbach, Franck Court, Tristan Bouschet

## Abstract

Organoids are powerful models of brain development and function. Mouse brain organoids reproduce the main steps of neurodevelopment *in vivo* and differentiate more rapidly than human organoids, making them scalable for screening applications. However, mouse brain organoids remain poorly defined. Here, we showed that the transcriptome of mouse brain organoids developed for three weeks is close to that of a neonatal mouse brain. Strikingly, organoids reproduced the majority of alternative splicing and polyadenylation site events that define a neural identity *in vivo*. Proteomics revealed that most changes in mRNA expression were translated into proteins. Mouse brain organoids differentiated for only three weeks already harboured a complex network of synaptic proteins, including the ionotropic and metabotropic receptors for the neurotransmitters glutamate and GABA. To conclude, our study provides compelling evidence that mouse brain organoids are a rapidly maturing and relevant model of mammalian brain development and function in a neonatal-like environment.

## Introduction

Since the landmark study by Yoshiki Sasai’s team describing the spontaneous self-organisation of dissociated mouse and human embryonic stem cells (ESCs) into brain-like tissues [1], brain organoids have emerged as a powerful tool for modelling brain development and function.

While human brain organoids capture specific features of human neurodevelopment, mouse brain organoids have several advantages over their human counterparts. First, brain organoids generated from mouse ESCs (mESCs) complete neuronogenesis in only three weeks, compared to several months for human organoids [2]. Mouse brain organoids recapitulate the main steps of neurogenesis that take place in most mammals [1,3,4]. The mouse is the predominant model for preclinical studies in the brain [5], and ESCs can be isolated from mouse lines [6,7]. Lastly, even if a study comparing their reproducibility is lacking, the emerging picture is that the generation of mouse brain organoids is more reproducible than human organoids, possibly because the building plan of the mouse is simpler and therefore easier to reproduce *in vitro*, and the limited size of mouse organoids facilitates oxygen and nutrient diffusion, thereby limiting necrosis.

Similar to human pluripotent cells, mESCs can be specified into organoids of defined brain identities, including the cerebral cortex and the cortical hem [1,3], the striatum [8], and the hypothalamus [9]. Mouse brain organoids recapitulate the cell diversity of the mouse brain, and they contain neural progenitors, neurons and glial cells [1,3,4], in the expected proportions [4]. Mouse brain organoids display spontaneous calcium waves [1] and electric activities resembling those recorded in embryonic brain slices [3], which indicates a certain degree of functional maturity.

In contrast to human brain organoids, whose transcriptome [10,11] and proteome [12,13] are well defined, the transcriptome of mouse brain organoids was only partly defined by single-cell RNA-seq [4], and their proteome is unknown.

Here, we have generated the transcriptome of mouse brain organoids at three developmental stages and showed that it resembled the mouse newborn brain. Beyond cell identity, we analysed their transcriptome to an unprecedented level and asked whether brain organoids recapitulate two key gene regulatory features critical for neurodevelopment: exon skipping [14] and alternative polyadenylation site usages [15]. Finally, brain organoids were profiled by proteomics to shed light on the relative abundance of proteins, including synaptic proteins, and to estimate whether changes in RNA levels are effectively translated into proteins.

## Results

### Transcriptomic profiling of mouse embryonic stem cells differentiated into brain organoids

mESCs were differentiated into brain organoids using a protocol [16] inspired by the original protocol developed by Sasai and co-workers [1], which consists in aggregating mESCs into ultra-low adhesion wells in a media supplemented with 10% Knock-out serum replacement (KSR), the BMP/Smad inhibitor DMH1 to promote neural induction and the Wnt inhibitor IWP-2 to rostralize the neural tube (**Fig. 1A**). After 7 days of culture, neural organoids were transferred to ultra-low adhesion plates in a maturation medium containing N2 and B27 without vitamin A [16], under orbital shaking to improve nutrients and oxygen transfers. The quality of differentiation was confirmed by immunofluorescence on starting mESC cultures (**Fig. 1B**) and cryosections on organoids developed for 7, 14, and 21 days **(Fig. 1C**). As expected, this protocol led to a massive expansion in organoid size and the generation of neural progenitors (NESTIN and PAX6^+^ cells), neurons (ELAVL3^+^, TBR1^+^, TUBB3^+^ cells) and astrocytes (GFAP^+^ cells) (**Fig. 1C**). Next, brain organoids (at days 7, 14, and 21) were profiled by RNA-seq. Since we and others [1,17,18] have shown that mESCs differentiated into brain cells for three weeks share analogies with the newborn brain (NBB), NBB samples were included as gold standards. In addition, we also profiled mESC samples, reasoning that because mESCs are derived from *in vivo* growing blastocysts [6,7], mESCs could serve as a reasonable starting developmental time point for both organoids and the NBB.

**Figure 1.**
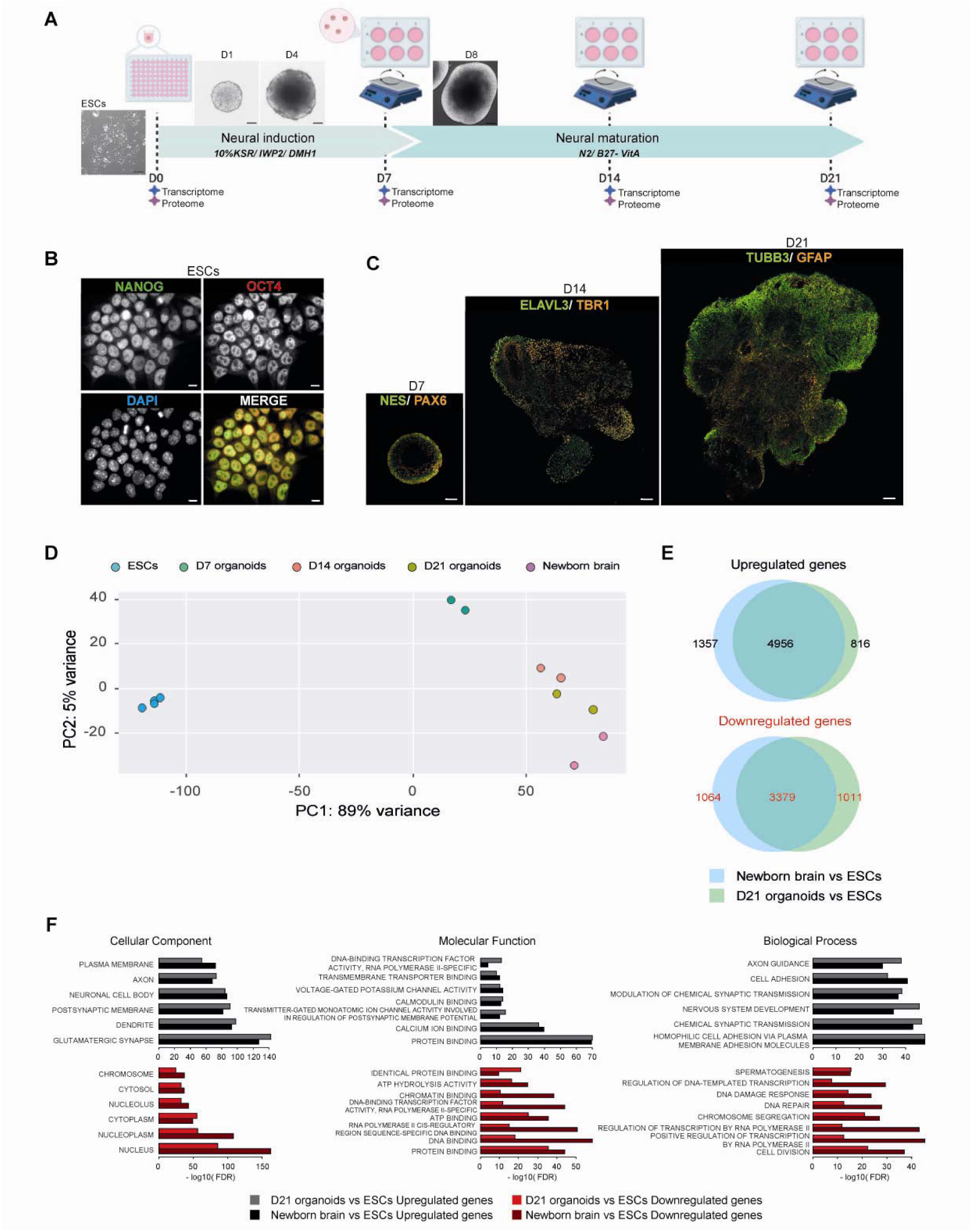
The transcriptome of mouse brain organoids progressively approaches that of the neonatal mouse brain. (A) Schematic timeline of brain organoids generation from mESCs. mESCs and brain organoids developed for 7, 14 and 21 days were profiled by transcriptomics and proteomics. Part of the figure was created with Biorender. (B) Immunostaining of mESCs-derived brain organoids. Cultures of mESCs were stained with NANOG (green), POU5F1/OCT4 (red), and nuclei with DAPI (blue). Scale bars: 10 µm. (C) Organoid cryosections were stained with NESTIN and PAX6 (at day 7), ELAVL3 and TBR1 (at day 14), and TUBB3 and GFAP (at day 21). Scale bars: 100 µm. (D) PCA showing the distribution of the transcriptomes of mESCs, brain organoids (at days 7, 14, and 21), and NBB samples. (E) Venn diagram showing the overlapping upregulated and downregulated genes between D21 organoids and NBB, both compared to mESCs. (F) Histograms of enriched GO terms in NBB vs ESCs and D21 organoids vs ESCs performed on upregulated genes (black and grey bars) and on downregulated genes (red and light red bars) determined using DAVID. For each term category (Cellular component, molecular function, and biological process), the five highest enriched terms in each gene list are shown.

### The transcriptome of mouse brain organoids progressively approached that of the neonatal mouse brain

To assess the similarity between brain organoids and the newborn brain, we compared their transcriptomes. Principal component analysis (PCA) showed that mESCs clustered separately from brain organoids and the NBB samples (**Fig. 1D**). Organoids at day 7 were the closest to mESCs (despite already a sharp transition in gene expression (**Fig. S1**), then D14 organoids, and D21 organoids were the closest to NBB samples. This suggests that the transcriptome of brain organoids progressively approaches that of the NBB. Most of the differentially expressed genes (DEGs) between D21 organoids and mESCs, and between the NBB and mESCs, were common, with 4956/7129 (69.5%) common genes for upregulated genes, and 3379/5454 (61.5%) for downregulated genes (**Fig. 1E**). Accordingly, brain organoids and NBB samples shared the same most significantly enriched Gene Ontology (GO) terms. For upregulated genes (**Fig. 1F**), this includes, for cellular component, ‘glutamatergic synapse’ (enrichment FDR: 2.59e-143 for brain organoid, and 6.26e-128 for the NBB), ‘dendrite’ (enrichment FDR: 1.86e-98 for brain organoid, and 2.34e-93 for the NBB), and ‘axon’ (enrichment FDR: 8.35e-74 for brain organoid, and 6.75e-69 for the NBB). Among common terms between brain organoids and the native brain (all with an enrichment FDR<1e-10), we also retrieved ‘transmitter-gated monoatomic ion channel activity involved in the regulation of postsynaptic membrane potential’ (molecular function) and ‘homophilic cell adhesion via plasma membrane adhesion molecules’ (biological process) (**Fig. 1F**). For downregulated genes (bottom panels of the **figure 1F**), the most significantly enriched terms were also common between organoids and the NBB and include ‘cell division’. Thus, GO analysis suggests that the transcriptome of D21 organoids is enriched in the same terms associated with brain function and development as the newborn brain.

### Brain organoids lacked the non-neural cell types present in the neonatal mouse brain

However, PCA (**Fig. 1D**), Venn diagrams (**Fig. 1E**), and a heatmap reporting gene expression across samples (**Fig. S2A**) indicated a partial difference between the transcriptome profiles of D21 brain organoids and NBBs. We identified 804 upregulated and 315 downregulated genes when comparing the NBB to D21 organoids (**Fig. S2 B-C**). As shown in **Figure S2D**, genes upregulated in the NBB were enriched in GO terms for ‘angiogenesis’, ‘regulation of blood pressure’, ‘inflammatory response’, ‘osteoclast differentiation’, and ‘extracellular matrix’. This likely reflects that, compared to the native brain, organoids lack blood cells, a circulatory system, bone cells (skull), and cells triggering an inflammatory response in the brain (principally microglia), all of which might participate in depositing a brain-specific matrix.

### Brain organoids largely reproduced alternative splicing events

Next, to extend our analysis of the transcriptomes of brain organoids and the NBB beyond cell identity, we asked whether gene regulatory features important for brain development are well reproduced by organoids. We first focused on alternative splicing, which is finely regulated during development and leads to splice isoforms that are tissue-specific [14]. However, it remains unknown to what extent brain organoids (including human organoids) recapitulate the variety of transcript isoforms found in the in vivo brain. To compare splicing events occurring in the NBB and D21 organoids (both compared to mESCs), we employed the widely used statistical suite R-MATS -Robust and flexible detection of differential alternative splicing from replicate RNA-Seq data [19]. During differentiation, we observed a profound remodelling of transcripts both *in vivo* and *in vitro* (**Fig. 2A**). Some genes were affected by several alternative splicing events (hence, the number of genes < number of events). Most of the changes in transcript isoforms related to exon skipping, which results in either exon inclusion or exclusion (**Fig. 2A**). Thus, we next focused on comparing the *in vivo* and *in vitro* situations for exon skipping. Strikingly, on 1,258 genes with skipped exons in the newborn brain, 968/1,258 (77%) were also present in D21 brain organoids (**Fig. 2B**). Interestingly, the number of skipped exon events increased during organoid development (**Fig. S3A**), and the correlation between the R-MATS metric of the NBB compared to organoids also increased as development progresses (**Fig. S3B**). Thus, collectively, this suggests that brain organoids progressively acquire part of the repertoire of the splice isoforms of the NBB.

**Figure 2.**
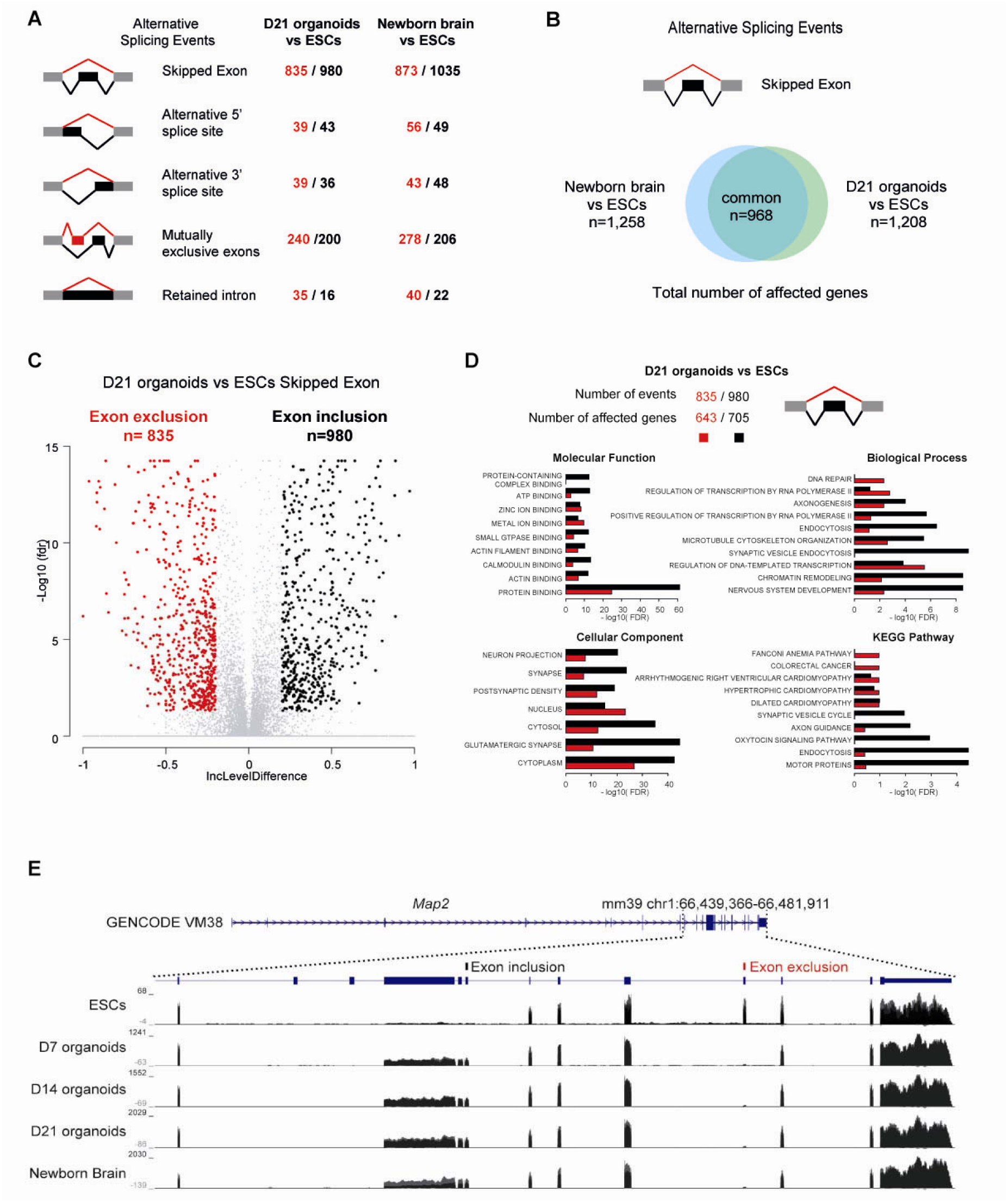
Brain organoids largely reproduce exon skipping events. (A) Alternative splicing events detected using R-MATS in the NBB and D21 organoids compared to mESCs. Red and black numbers correspond to the nature of the alternative splicing events (red: exon exclusion; black: exon inclusion). (B) Venn diagram showing the number of overlapping genes affected by skipped exon events between brain organoids at day 21, and the NBB, both compared to mESCs. (C) Volcano plot of exon skipping events in D21 organoids compared to mESCs. There were 835 events of exon exclusion (red) and 980 events of exon inclusion (black). Alternative splicing events were identified when |IncLevelDifference| >0.2 and FDR <0.05. (D) Histogram of GO analysis (KEGG pathway, cellular component, molecular function, and biological process) of genes affected by exon exclusion (red) or inclusion (black) in D21 organoids compared to mESCs. (E) UCSC genome browser capture of RNA-seq data of mESCs, brain organoids (D7, D14, and D21), and NBB for reads mapping to the *Map2* gene. The genomic location of the excluded exon is indicated by vertical red bars and inclusion by black bars.

Next, we performed a GO analysis on the genes displaying either exon inclusion or exclusion in D21 organoids compared to mESCs (**Fig. 2C**). GO analysis revealed an enrichment in neural terms, including ‘axonogenesis’, ‘axon guidance’, ‘synaptic vesicle’, ‘neuron projection’, and ‘glutamatergic synapse’, independently on the nature of events (exon inclusion or exclusion) (**Fig. 2D**). Among the list of genes that were common between the NBB and D21 organoids, we noticed the presence of *Map2*, a gene involved in neurite outgrowth, synaptic plasticity, and transport [20]. A screen capture of RNA-seq data uploaded into the UCSC genome browser confirmed that compared to the ESC stage, *Map2* experienced both exon inclusion and exon exclusion, leading to similar *Map2* transcript(s) isoforms in the organoids and the newborn brain (**Fig. 2E**). Thus, collectively, these data suggest that alternative splicing events, notably exon skipping, is well reproduced by brain organoids and that is associated with a neuronal signature.

### Brain organoids largely reproduced alternative polyadenylation events

Another gene regulatory feature important for neurodevelopment is alternative polyadenylation (APA), which results in either 3’UTR lengthening or shortening [15]. APA is context-dependent and important for transcript location, stability and translation [15]. In the developing brain, a defined subset of transcripts is targeted by APA, which most often results in 3’UTR lengthening [21,22]. However, it remains elusive whether brain organoids recapitulate the APA site usages of the *in vivo* brain. Thus, we next sought to identify genes experiencing APA in brain organoids and compare that list to the *in vivo* brain. To this aim, we employed the bioinformatics package APAlyzer [23]. When compared to the ESC stage, we identified 307 genes with APA in organoids at day 7 (**Fig. S4A**), 360 genes in organoids at day 14 (**Fig. S4B**), 427 genes in organoids at day 21 (**Fig. 3A**), and 296 genes for the NBB (**Fig. 3B**). For all samples, the majority of APA resulted in 3’UTR lengthening (**Fig. 3A**, **B** and **Fig. S4**), as previously reported for the mouse brain [21,22]. More specifically, in brain organoids at day 21, 366 transcripts lengthened while 61 shortened and for the NBB, 258 transcripts extended and 38 shortened (**Fig. 3A**). Remarkably, 171/258 (66%) transcripts that extended in the NBB also extended in D21 brain organoids (**Fig. 3C**). An APA usage heatmap confirmed that D21 organoids globally mimic the NBB situation (**Fig. 3D**). Among the genes identified by APAlyzer, we noticed the presence of the RNA-binding protein *Elavl1*, known to be longer in the brain [24] and the scaffolding protein *Homer2*. The extension of the 3’UTR of *Elavl1* and *Homer2* was confirmed by looking at RNA-seq data (**Fig. 3E**, **F**). The 366 genes whose transcripts lengthen in D21 organoids were enriched in GO terms related to the ubiquitin-proteasome pathway, including ‘Protein Ubiquitination’ (Biological process), ‘Ubiquitin protein ligase binding ‘(Molecular function), and ‘Ubiquitin mediated proteolysis’ (KEGG pathway) (**Fig. S5**). Thus, brain organoids recapitulated most of the APA events taking place *in vivo*. This resulted in 3’UTR lengthening over shortening and in an enrichment in genes related to the ubiquitin proteasome pathway.

**Figure 3.**
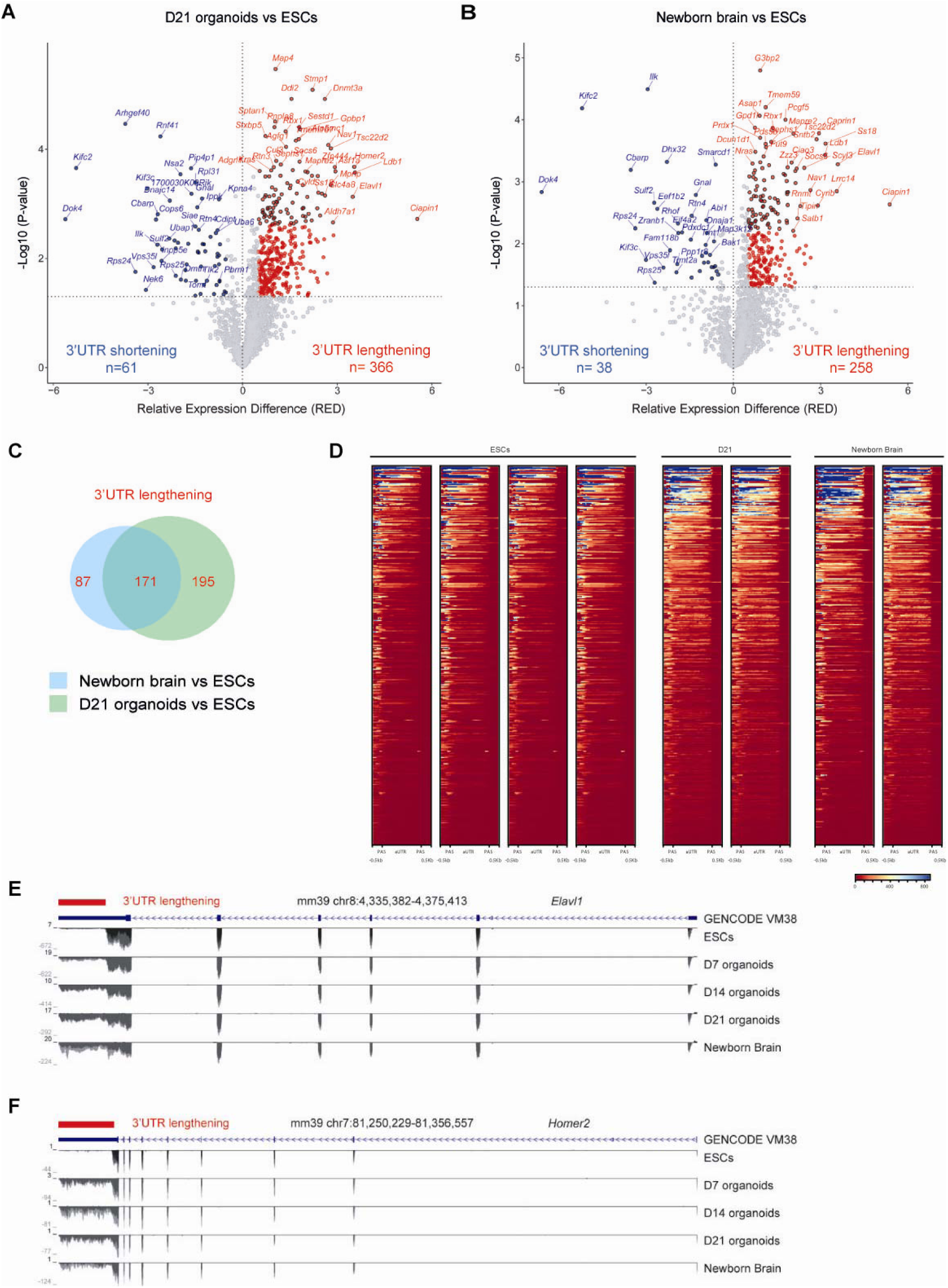
Brain organoids largely reproduce alternative polyadenylation site usages. (A) Volcano plot of the distribution of genes identified with alternative polyadenylation site usage in D21 organoids compared to mESCs, indicating whether a proximal (blue, resulting in 3’UTR shortening) or a distal (red, resulting in 5’UTR shortening) site was used. Genes were identified using APA analyser with the following cut-offs: |RED| (Relative Expression Difference) between proximal and distal polyadenylation site usage>0.5 and p-value <0.05. (B) Same as in A, but for the NBB compared to mESCs. (C) Venn diagram showing the number of overlapping genes displaying 3’UTR lengthening in NBB and D21 organoids. (D) Heatmap showing APA lengthening for genes identified comparing ESCs and D21 organoids in RNAseq of ESCs, D21 organoids and NBB samples. (E) UCSC genome browser capture of RNA-seq data of mESC, brain organoids (D7, D14, and D21), and NBB for reads mapping to the *Elavl1* gene. The genomic location of the site with APA usage is indicated by horizontal red bars. (F) Same as in E, but for the *Homer2* gene.

### Deciphering the proteome of mouse brain organoids

While RNA-seq is widely used on human brain organoids [10,11], proteomics studies are still rare [12,13], and only one study compares the transcriptome and proteome of human brain organoids [25]. In addition, the proteome of mouse brain organoids is unknown. Thus, we next sought to characterise the proteome signature of mouse brain organoids at days 7, 14, and 21 and compare them to their transcriptome. To this aim, mESC (n=4 samples), and brain organoids at day 7 (n=3), day 14 (n=4), and day 21 (n=4) were analysed by Liquid Chromatography coupled to tandem Mass Spectrometry -LC-MSMS-, performed on an Orbitrap device (see *Methods*). This approach yielded the identification of 5901 proteins with a cut-off of> 3 peptides per protein. In PCA, the ESC and D21 organoid samples clustered apart with D7 and D14 organoids in between them (**Fig. 4A**), as observed for RNA-seq (**Fig. 1D**). Accordingly, a heatmap representing variation in protein expression across samples revealed massive changes in protein expression during the differentiation process, in particular during neural induction (**Fig. 4B**). Next, we identified differentially expressed proteins (logFC > 2, and p>0.01) during the transition from mESC to D7 organoids (**Fig. 4C**), from D7 to D14 organoids (**Fig. 4D**), and from D14 to D21 organoids (**Fig. 4E**). As suggested by the distance between the four developmental stages in the PCA, more proteins were differentially expressed between ESC and day 7 and between day 7 and day 14 (430 proteins) than between d14 and day 21. This indicates that there are sharp transitions in the proteome signature from ESC to day 7 and from day 7 to day 14, while the proteome is relatively stable between day 14 and day 21. As expected, we confirmed the selective expression of proteins that mark cell identity: the stemness marker NANOG was enriched in mESCs and the neural progenitor marker NESTIN in D7 organoids (**Fig. 4C**). The radial glia FAPBP7 and the axonal marker MAPT were enriched at day 14 -compared to day 7- (**Fig. 4D**). D21 organoids were enriched (compared to D14 organoids) in the mature neuronal markers THY1, GRM5 and CAMK2B, in the oligodendrocyte marker CNP and in the astrocyte marker GFAP (**Fig. 4E**).

**Figure 4.**
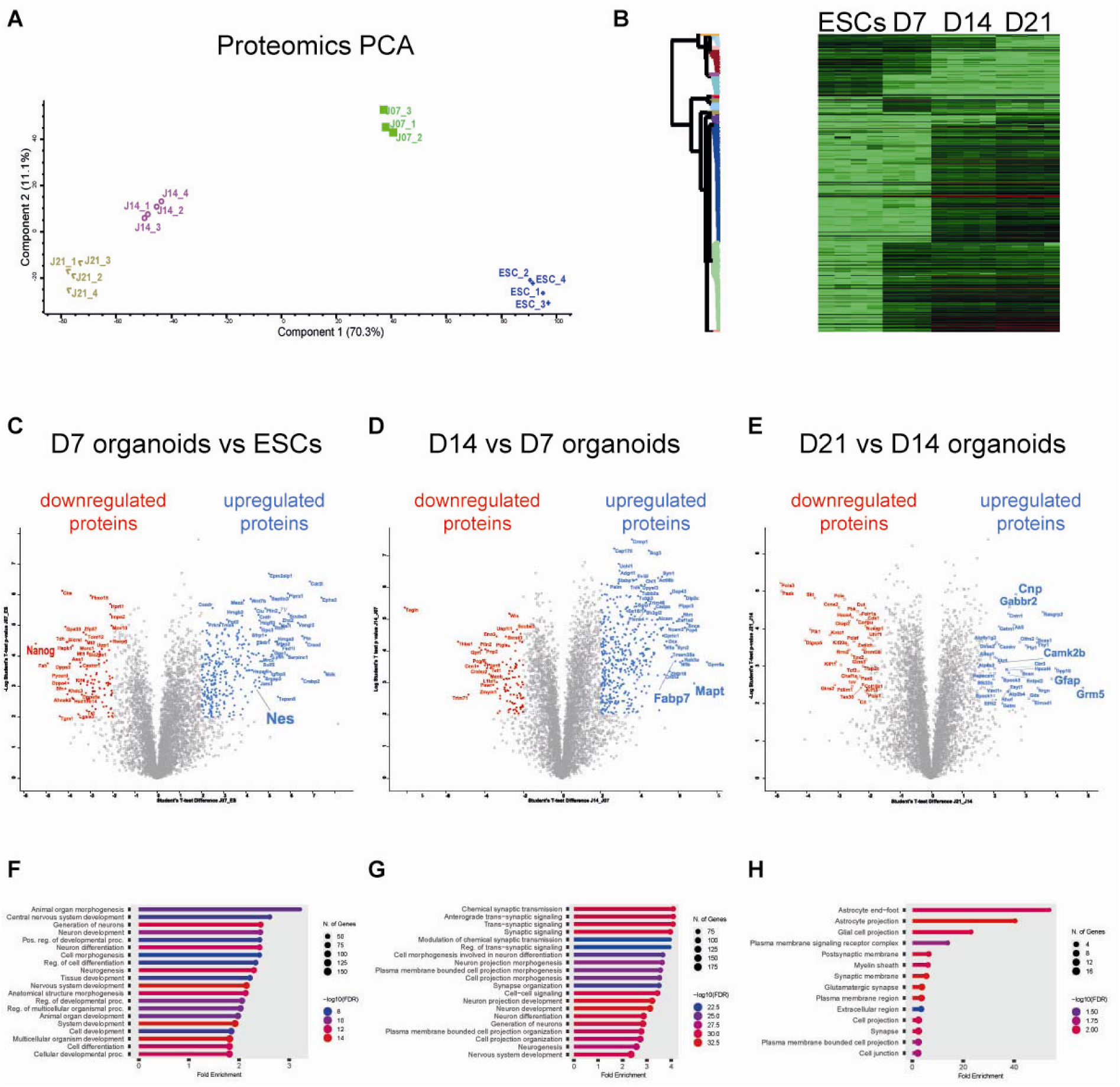
The proteome of developing brain organoids. (A) PCA showing the distribution of the proteomes of mESCs and brain organoids (at days 7, 14, and 21). (B) Heatmap of the relative expression of proteins across samples. (C-E) Volcano plots of differentially expressed proteins between D7 organoids and mESC (C), D14 and D7 organoids (D), and D21 and D14 organoids (E). Proteins significantly downregulated are indicated in red, and upregulated proteins are in blue. A selection of cell fate markers is written in a larger font. (F) Histograms of enriched GO biological processes terms in D7 organoids vs ESCs determined using Shiny GO. (G) Histograms of enriched GO biological processes terms in D14 organoids vs D7 organoids. (H) Histograms of enriched GO Cellular component terms in D21 organoids vs D14 organoids.

Next, to infer the identity of organoids via their proteome, we performed a GO analysis on differentially expressed proteins across the three developmental transitions. GO analysis was performed using Shiny GO, which contains a large database derived from Ensembl and STRING-db [26]. In organoids at day 7 (compared to mESCs), the top enriched pathways included ‘multicellular organism development’ (enrichment FDR: 6.9E-15), ‘central nervous system development’ (enrichment FDR: 4.0E-09), and ‘neuron differentiation’ (enrichment FDR: 8.4E-13) (**Fig. 4F**). Strikingly, the proteins enriched in D14 organoids compared to D7 organoids indicated an additional sharp transition towards maturity with terms such as ‘chemical synaptic transmission’ (enrichment FDR: 1.1E-31) and ‘neuron projection development’ (enrichment FDR: 4.0E-33) (**Fig. 4G**). In D21 organoids compared to D14, there was no enrichment in biological process terms (not shown). However, proteins were enriched for cellular component terms, suggesting that organoids continued maturing (‘Glutamatergic synapse’, enrichment FDR: 6.0E-03; ‘Myelin sheath’: 1.1E-02, and ‘astrocyte end-foot’, 1.1E-02) (**Fig. 4H**). Thus, these experiments suggest that the proteome drastically evolved between mESC and D7 organoids and between D7 and D14 organoids, and then slightly evolved.

### The proteome of brain organoids encompasses morphogens, cortical-layer markers and chemo-attractive and repulsive cues

Morphogens shape the developing brain (Sur and Rubenstein 2005; O’Leary et al. 2007). Brain organoids lack the organisers that secrete morphogens *in vivo*; accordingly, the identity of organoids is triggered by mimicking the influence of organisers through the addition of exogenous morphogens [29] or by including morphogen centres [30]. Of interest, our model of organoids mimics the development of the most anterior part of the neural tube, which develops in the absence of morphogens [1]. However, we cannot rule out that these organoids generated ‘by default’ produce their own morphogens. The identity of secreted morphogens (if any) by organoids is elusive. Thus, we next sought to determine the repertoire of morphogens, growth factors and cognate receptors intrinsically expressed by brain organoids. As shown in **Figure 5A**, we detected WNT7B and WNT8B proteins (together with the Wnt receptors FZD2 and FZD7), which all peaked at day 7. BMP1 and BMPR2 were detected at later time points, consistent with a role in astrocyte generation and maintenance [31]. We did not detect SHH, any FGFs, R SPONDINs, or their receptors (SMO, FGFRs, and LGR5, respectively). The EGFR, important for the fate of neural stem cells [32], was also expressed, but not the EGF. One question is, therefore, what drives the growth of organoids? During neural induction, the only exogenous growth factor protein is insulin, present in the KSR [9], and we detected the insulin receptor (INSR) in all developmental stages (although it was not detected in all samples). Thus, insulin transducing via the INSR might be the primary signalling pathway responsible for organoid growth. Insulin might work in conjunction with lipids since the KSR contains lipid-rich albumin [33], and we found that brain organoids expressed the apoE/lipoprotein receptor LRP, known to mediate lipid uptake in the brain [34].

**Figure 5.**
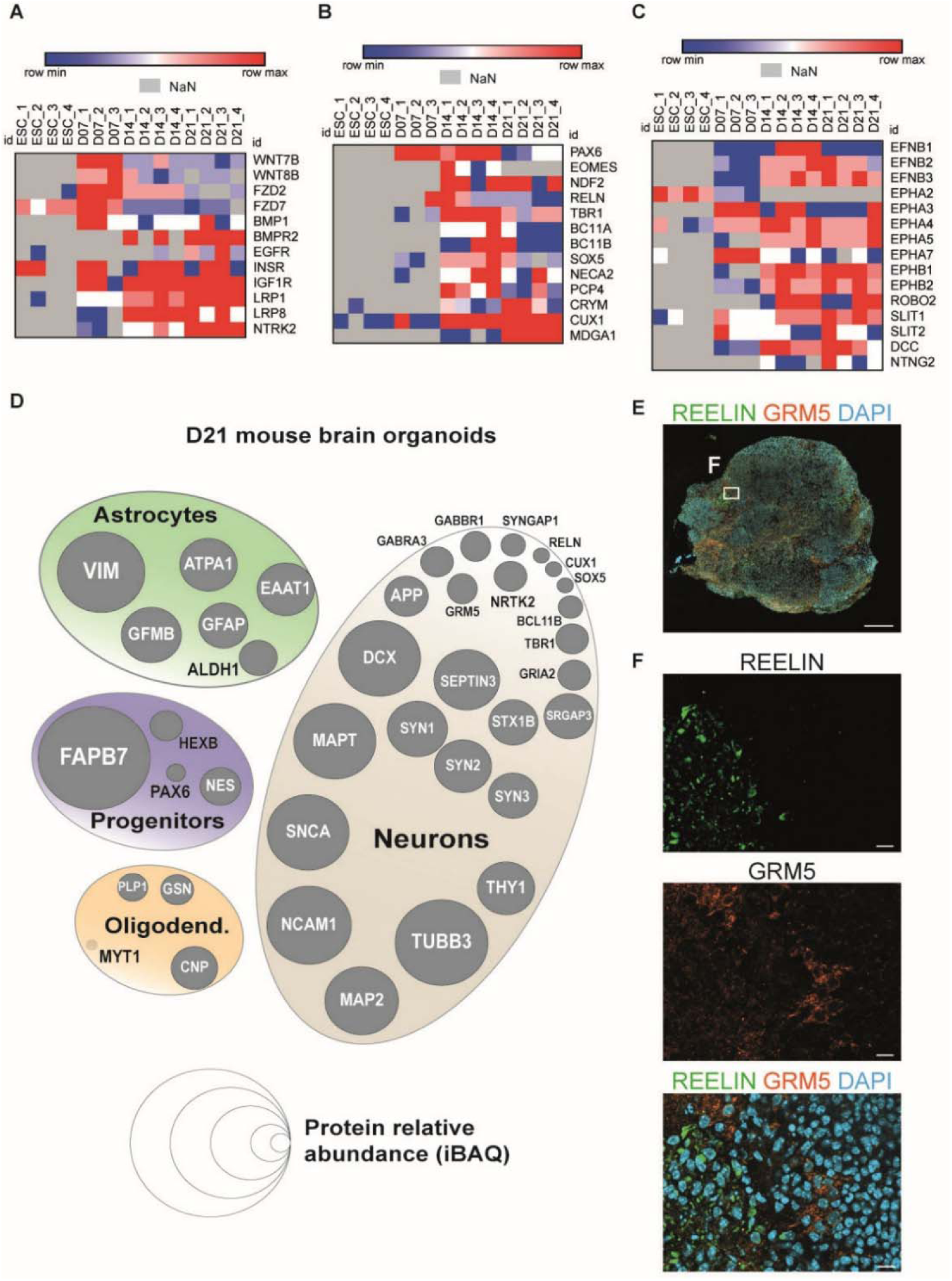
Evolution of the proteome during the development of brain organoids and the relative abundance of known brain proteins in D21 organoids. (A) Heatmaps reporting the level (low: blue, average: white, high: red) of protein expression across samples for a selection of morphogens, growth factors and cognate receptors. The heatmap was generated using Morpheus (the Broad Institute, MIT). ‘Not a Number’ (NaN) values are in grey. (B) Heatmap of protein expression for cortical layer markers. (C) Heatmap of protein expression for chemoattractive/chemorepulsive molecules. (D) Proteins known to be expressed in neural progenitors, neurons, astrocytes and oligodendrocytes are represented as concentric circles proportional to their relative abundance (iBAQ) in D21 organoids. (E) D21 organoid cryosections were stained with REELIN (green), GRM5 (orange), and DAPI (blue) and imaged at a fluorescent microscope using the apotome mode. Scale bar: 200 µm. (F) High magnification images from the inset drawn in E. Scale bars: 10 µm.

In the developing cortex, stem cells integrate a combination of morphogens, growth factors and intrinsic cues to produce cells in a timely, invariant order: PAX6+ neural progenitors are generated first, followed by EOMES/TBR2+ progenitors. Next, these progenitors progressively generate neurons of deep, then upper-layer identities [35,36]. As shown in **Figure 5B**, the timing of appearance of protein markers of cell identity in brain organoids reflected quite well the *in vivo* situation: PAX6 appeared first (from day 7), EOMES/TBR2 was exclusively expressed at day 14, and, for neurons, REELIN (a marker of Cajal-Retzius neurons) appeared first, followed by TBR1 (layer VI), deep layers (BCL11A/B and SOX5), and upper layers (CRYM and MDGA1). This suggests that the temporal specification of corticogenesis is quite well maintained in mouse brain organoids.

In the developing brain, neurons acquire a complex repertoire of chemoattractive/chemorepulsive molecules, including EPHRIN receptors and ligands [37] and SLIT/ROBO [38]. These intercellular communication systems position cells, delimitate tissue boundaries and guide navigating axons towards their targets. We found that brain organoids expressed the transmembrane EPHRINS B1-B3, but not the GPI-anchored Type A EPHRINS, as well as some EPHA and EPHB receptors (**Fig. 5C**). As reported by Fernandez-Alsonso and colleagues [39], we observed that EPHA2 was enriched in pluripotent cells (**Fig. 5C**). ROBO2 and SLIT1 and their regulators NETRIN and DCC were also detected by proteomics (**Fig. 5C**). Thus, collectively, brain organoids harbour a complex repertoire of proteins known to be involved in neurodevelopment.

### Determination of the relative abundance of brain-specific proteins in brain organoids

Next, we sought to determine the relative abundance of proteins in brain organoids. To this aim, we took advantage of intensity-based absolute quantification (iBAQ) values [40]. As previously described for human brain organoids [12], we represented the abundance of proteins known to be selectively enriched in defined brain cell types as circles whose size are proportional to their iBAQ values (**Fig. 5D**). We focused on day 21 organoids and detected an abundant expression of the radial glia marker FABP7 (median iBAQ value:1,11E+09) while the pan neural progenitor marker NESTIN was less expressed (median iBAQ value:5,54E+06). This supports the view that in D21 organoids, the most represented type of neural progenitors is radial glia, the cell type that generates neurons [41]. The proteome of D21 brain organoids also contained the astrocyte markers VIMENTIN (median iBAQ value: 1.83E+08) and GFAP (8,51E+06) (**Fig. 5D**). There was also an abundant expression of proteins known to be enriched in neurons, including cytoskeletal proteins (TUBB3; median iBAQ value: 4,00E+08, MAP2; 7,54E+07, and MAPT: 1,19E+08). We also detected proteins involved in synaptic transmission, including SYNAPSINs 1, 2, and 3, the ionotropic and metabotropic receptors for glutamate (GRM5 and GRIA2, respectively) and for GABA (GABBR1 and GABRA3, respectively) (**Fig. 5D**). The neurotrophin NRTK2/TRKB receptor, which regulates synaptic strength [42], was also expressed. Not surprisingly, and as indicated by the relative size of circles, all of these receptors were less abundant than neuronal filaments. We also detected markers of layers of the cerebral cortex [17,35]: REELIN for Cajal-Retzius neurons; TBR1, SOX5, and BCL11B for deep layers, and CUX1 for upper-layer neurons (**Fig. 5D**). As predicted by proteomics, we confirmed that both REELIN and GRM5 were expressed in brain organoids (**Fig. 5E**). The typical markers of the oligodendrocyte lineage, PLP1 (median IBAQ value:1,84E+06), GSN (2.31E+06), and CNP (7.47E+06) were less abundant than those of radial glia, neurons, and astrocytes, indicating an underrepresentation of this cell type (**Fig. 5D**). We did not detect proteins known to be enriched in microglia (IBA1) and blood vessels (CD31/PECAM1). Collectively, this suggests that D21 organoids contain proteins enriched in the different cell types composing the newborn brain, except for microglia and blood vessels. In addition, the proteome of mouse brain organoids differentiated for three weeks already contains a complex repertoire of proteins involved in synaptic transmission.

### High correlation between differentially expressed RNAs and proteins during developmental transitions

Finally, we sought to determine whether changes in RNA expression are effectively converted into proteins. First, we noticed that out of the 15,939 gene products quantified by RNA-seq, 13,176/ 15,939 (82.6%) were protein-coding (**Fig. 6A**). On the other hand, proteomics detected the high number of 5,971 proteins (**Fig. 6B**). We reasoned that since changes in RNA and proteins over developmental time points were both quantified by a log2-fold change, we could directly compare variations in RNA and proteins using a Pearson correlation. In all three developmental transitions (from ESCs to D7, D7 to D14, and D14 to D21), the Pearson coefficient r >0.65 (**Fig. 6D-F**), indicating a highly positive correlation between protein and RNA variation during the genesis of mouse brain organoids. Visualising the changes in RNA and protein expression for a series of genes involved in development confirmed the parallel evolution of RNAs and proteins during organoid genesis (**Fig. 6G**, **H**).

**Figure 6.**
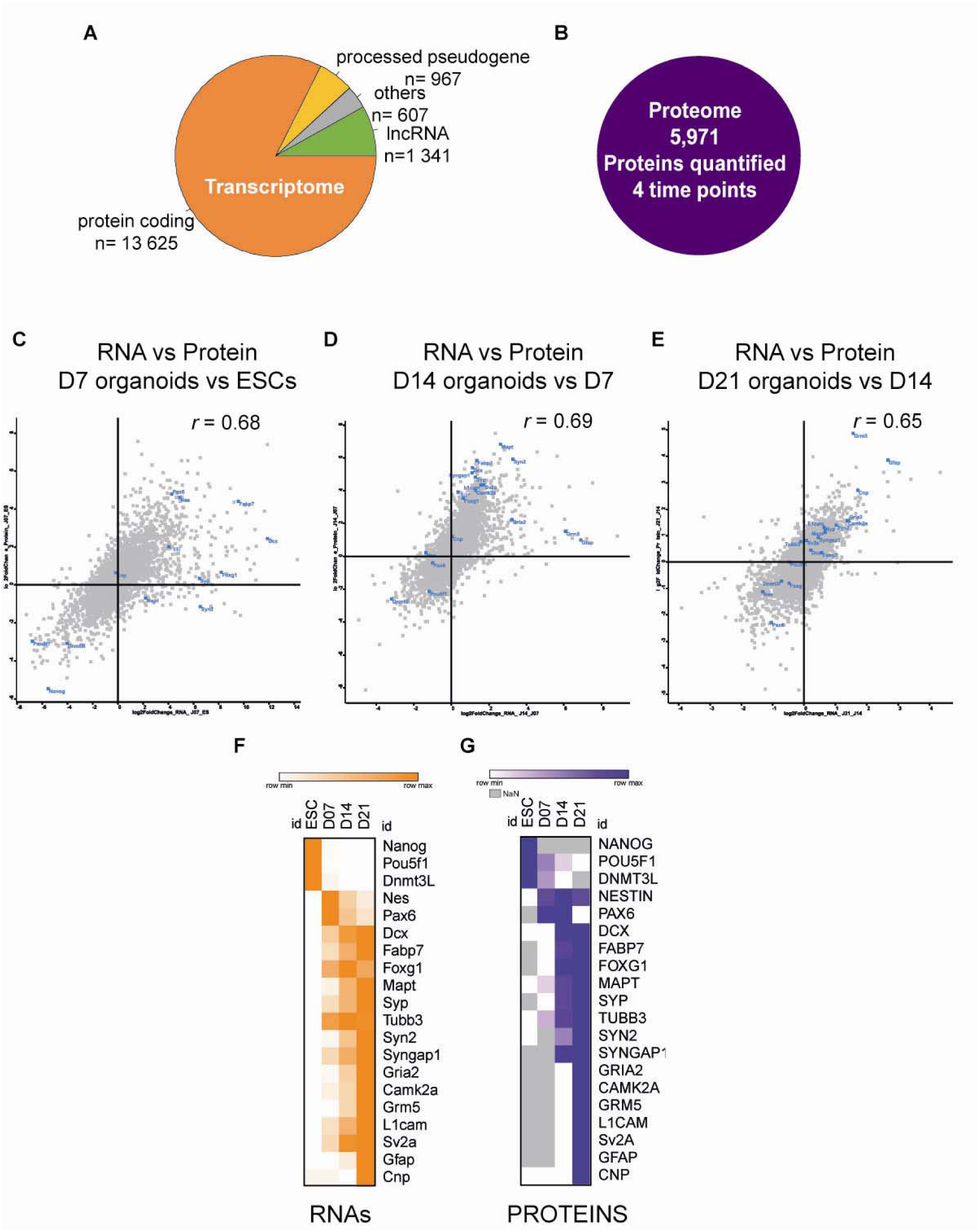
High correlation between differentially expressed RNAs and proteins during developmental transitions in brain organoids. (A) Gene annotation of the 16540 quantified genes with an average expression >1 FPKM in at least one group of samples. (B) Output of proteomics: 5971 proteins were quantified. (C, D, E) Log2-Fold changes in protein (Y-axis) and gene expression (X-axis) were plotted during developmental transitions from mESC to D7 organoids (C), from D7 to D14 organoids (D), and from D14 to D21 organoids (E), and Pearson coefficients r were calculated. (F) Heatmap reporting the levels (low: white, high: orange) of RNA expression across samples for cell fate markers. The heatmap was generated using Morpheus (the Broad Institute, MIT). (G) Heatmap reporting the levels (low: white, high: purple) of protein expression across samples for cell fate markers. The heatmap was generated using Morpheus (the Broad Institute, MIT). Not a Number (NaN) values are in grey.

## Discussion

Many brain diseases, including those that clinically manifest at adult age, such as Huntington’s and Alzheimer’s diseases, have a neurodevelopmental origin [43,44]. During the last century, insights into normal and pathological neurodevelopment have been gained from a variety of biological models, including C. elegans, Drosophila, mouse, and human samples. In this study, we present compelling evidence that mouse brain organoids are an attractive alternative model.

*In vitro* models that differentiate embryonic stem cells into brain cells in Petri dishes (in two dimensions, 2D) reproduce some cardinal features of the *in vivo* brain, including the sequential generation of cortical layers [17,45]. These models are also useful to discover genes involved in neurodevelopment [46] and human specification [47]. Concomitant with these 2D models, the Sasai group developed brain organoids [1], which were later improved by other groups to study microcephaly [48,49], and further complexified into brain assembloids [50].

Brain organoids have been generated from pluripotent cells of several species, including mouse [1], chimpanzee and macaques [51], and human [1,48,51]. The developmental timing of the brain generated from pluripotent cells is shorter for the mouse compared to primates [52]. Brain organoids capture these species differences [3,51]. The reproduction of human-specific features (including a complex genetic background) likely explains why most studies are being conducted on human organoids. However, mouse organoids represent an alternative model to study neurodevelopment and function because of their shorter generation time than human organoids. In addition, similarly to human cells, mouse ESCs can be specified into region-specific brain organoids [1,3,9], and they reproduce the cardinal features of *in vivo* development, including the sequential generation of cortical layers [1].

One likely advantage of mice over humans is their defined genetic background. By crossing C57Bl/6J laboratory mice to wild-derived inbred strains to generate F1 hybrid epiblast stem cells, Medina-Cano et al. have revealed the cis-acting transcriptional regulatory changes in mouse brain organoids [4]. Here, we have added a supplemental layer to our understanding of gene regulation in mouse brain organoids by showing that the transcriptome of mouse brain organoids progressively approaches that of the mouse newborn brain. Gene ontology analysis revealed an enrichment in the same most significantly enriched neural terms as the NBB, while an absence of non-neural terms. This contrasts with our previous work on *in vitro* cortex derived from mESCs in a Petri dish, where we noticed an expression of endoderm and mesoderm genes, suggesting the presence of (unwanted) cell types of other developmental origins in the 2D model [18].

Here, we have further observed that mouse brain organoids robustly reproduced two features of gene regulation important for setting a neural identity: alternative splicing and polyadenylation site usage [15,14], which, overall, resulted in exon skipping and 3’UTR lengthening. Thus, brain organoids might represent an attractive model to study these important gene regulatory mechanisms [15,14]. Genes whose transcripts experienced exon alternative usage (whether it was an exon inclusion or exclusion) were related to neurodevelopment. More unexpectedly, for extending transcripts, in addition to genes known to play a role at the synapse, such as *Homer2*, we found an enrichment for the ubiquitin-proteasome pathway. The ubiquitin-proteasome pathway is known to be essential for synaptic plasticity by maintaining protein homeostasis via protein clearance [53], but whether the act of 3’UTR lengthening plays a role remains to be defined.

We also observed that during the generation of brain organoids, the majority of changes in RNA levels are translated into changes at the protein level. We detected a high number of proteins (5971, all stages included) and observed a profound remodelling of the protein landscape during developmental transitions from pluripotency to neural induction (D7 organoids), and during the neural maturation of organoids (from day 7 to day 21). Notably, the proteome of brain organoids at day 14 indicates a certain degree of maturity, in agreement with calcium recordings performed on mouse brain organoids generated according to a ressembling protocol [1]. Ciarpella and co-workers [54] also reported the generation of brain organoids from mouse neural stem cells and evidenced a functional network maturation after less than 5 weeks.

Brain organoids also expressed a selection of proteins of the Ephrins/Ephrin-receptors and Slit-Robo families, which mediate intercellular communication. This finding might explain why, upon grafting, the axons of transplanted neurons generated *in vitro* from pluripotent stem cells navigate and connect with the appropriate targets in the host brain [17,45,55,56]. However, the combination of Ephrins/Ephrin-receptors/ Slit-Robo is cell type-specific, and we performed bulk proteomic experiments on brain organoids that contain a myriad of cell types. Single-cell proteomics could serve to establish whether the combination of Ephrins/Ephrin-receptors/ Slit-Robo in brain organoids is cell type-specific.

Finally, thanks to iUBAQ values, we have determined the relative abundance of proteins and uncovered that most of the synaptic components are present in organoids at day 21. In contrast to the proteome of human brain organoids, which did not contain REELIN [12], a protein important for neuronal migration in the cerebral cortex [57], REELIN was found in the proteome of mouse brain organoids. We also detected the presence of the REELIN receptor LRP8/ApoER2, as well as proteins known to be enriched in deep and upper-layer cortical neurons (TBR1, BCL11B, SOX5, and CUX1).

To conclude, despite the lack of important brain cell types from other developmental origins (blood vessels and microglia) that participate in brain development and function *in vivo*, an absence observed in all brain organoids so far, our study indicates that mouse brain organoids represent a relevant model to study neurodevelopment in mammals, particularly gene regulation and brain-enriched receptors such as GABAergic and glutamatergic receptors in a neonatal-like environment.

## Supplementary material

**Figure S1.**
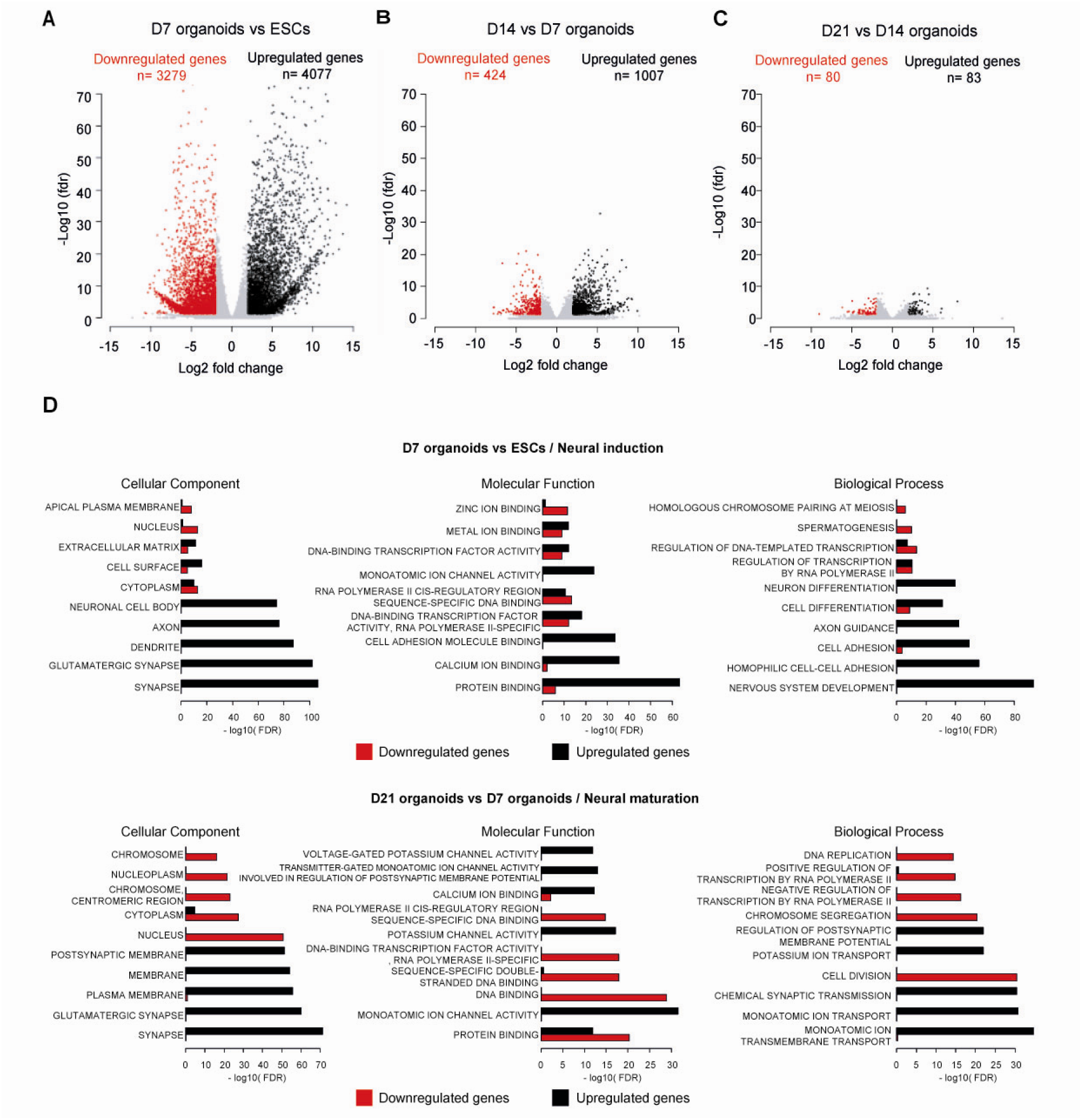
Functional enrichment analyses of differentially expressed genes during neural induction and maturation. (A-C) Volcano plot of the differentially expressed genes between D7 organoids and mESCs (A), D14 organoids and D7 organoids (B), D21 organoids and D14 organoids (C), illustrating the neural induction and the neural maturation of the brain organoids generation. The red and black dots respectively represent genes that were significantly down or upregulated during the two developmental stages (|log2FoldChange| >2, FDR <0.05). (D) Gene Ontology (GO) terms enriched for differentially expressed genes during neural induction (D7 organoids vs ESC) and neural maturation (D21 vs D7 organoids). For each GO term category, the five highest terms are shown for the differentially expressed gene lists. Red and black bars represent the –log10(FDR) of respectively the downregulated and upregulated gene lists.

**Figure S2.**
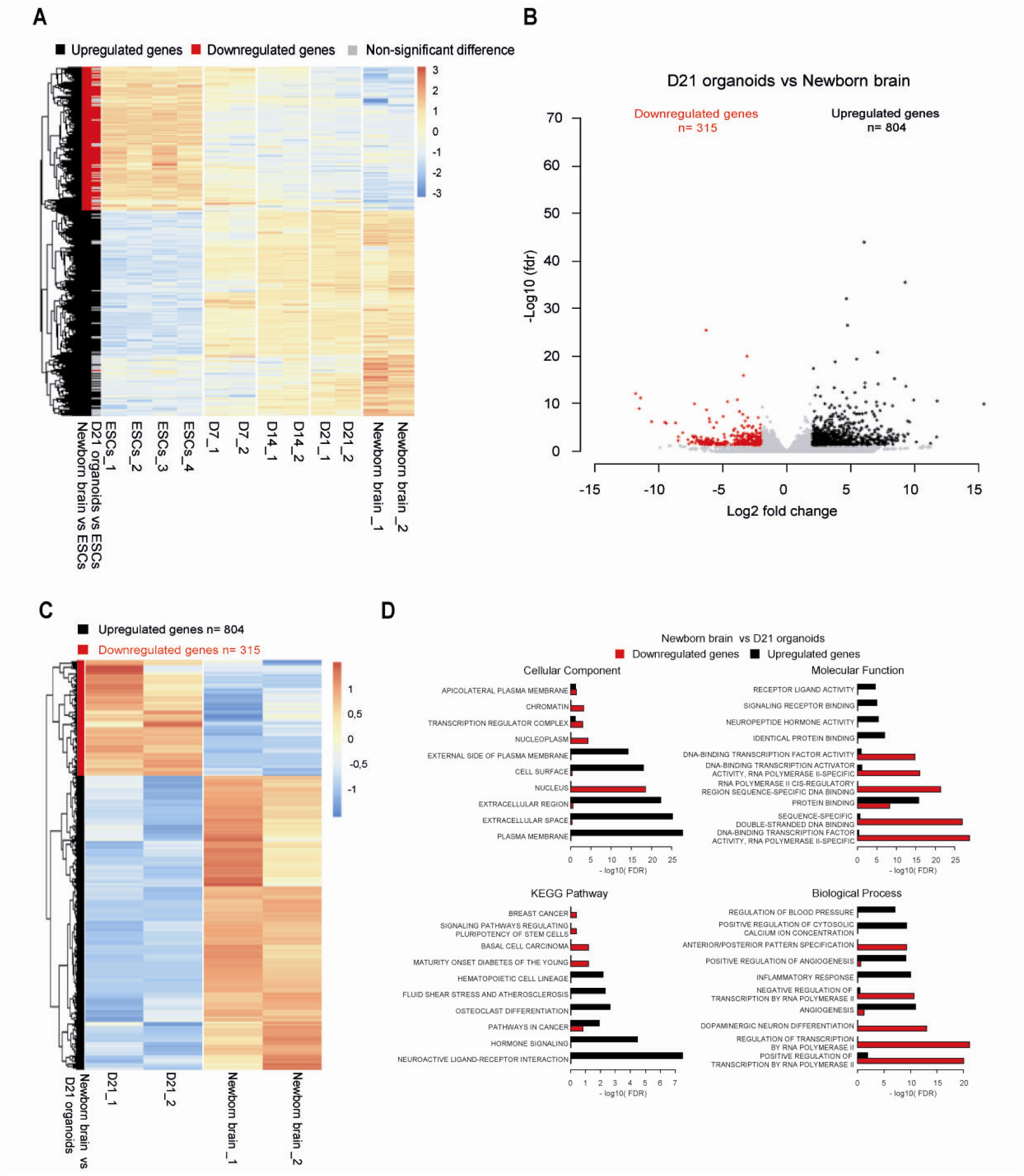
The transcriptome of mouse brain organoids progressively approaches the newborn brain. (A) Heatmap showing the centred-scaled expression levels of the differentially expressed genes between newborn brain and mESCs for all samples. Results of the differential analyses are shown on the left of the heatmap with black for upregulated genes, red for downregulated genes, and grey for non-significant differences. (B) Volcano plot of the differentially expressed genes between D21 organoids and newborn brain. Brain organoids generation was associated with the underexpression of 315 genes (red) and the overexpression of 804 genes (black) (C) Heatmap showing the centred-scaled expression levels of the differentially expressed genes between newborn brain and D21 organoids. Results of the differential analyses are shown on the left of the heatmap, with black for upregulated genes and red for downregulated genes. (D) Functional enrichment analyses of differentially expressed genes between newborn brain and D21 organoids. For each term category, the five highest terms are shown. Red and black bars represent the –log10(FDR) of respectively the downregulated and upregulated gene lists.

**Figure S3.**
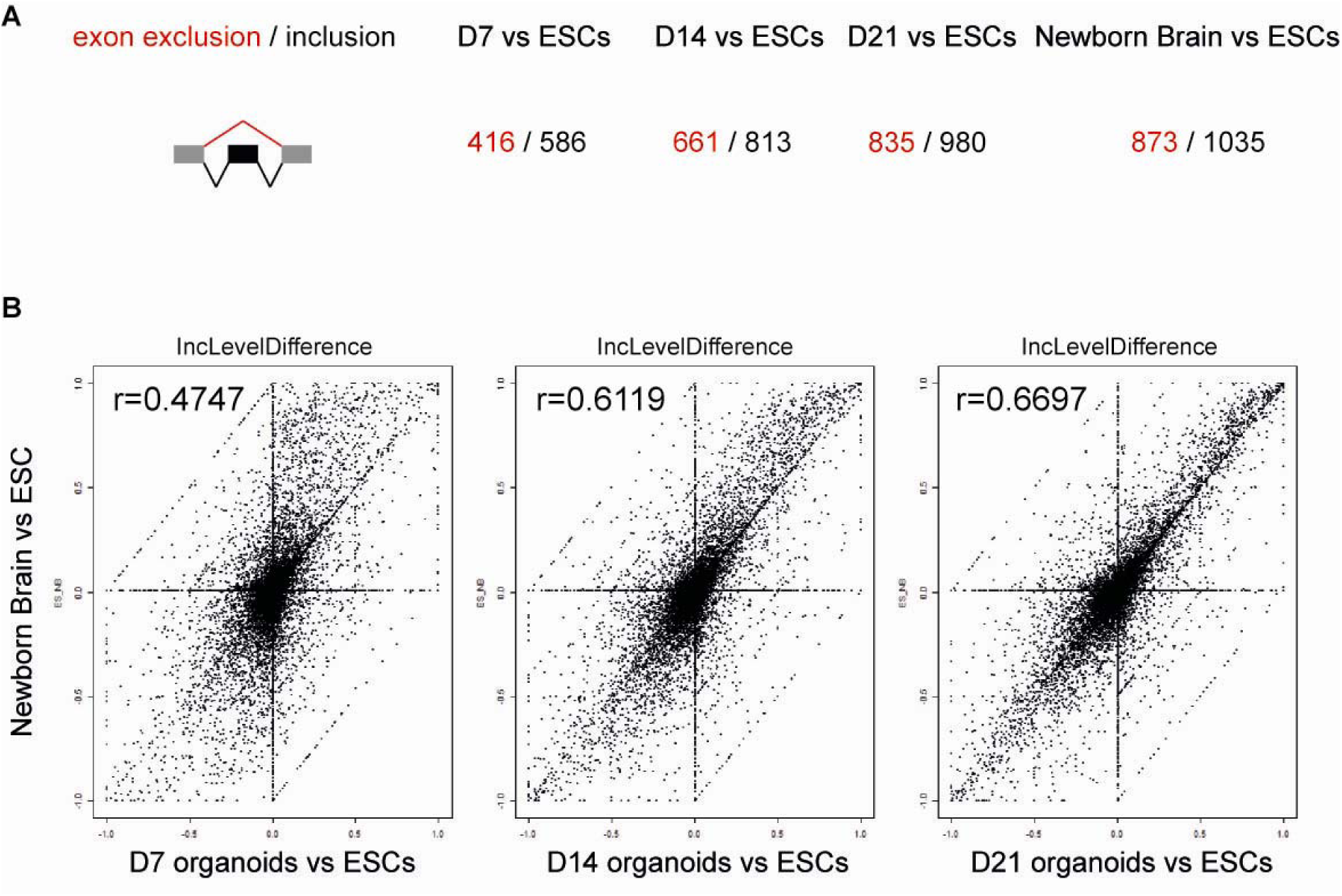
Evolution of alternative exon usage during organoid formation. (A) Evolution of alternative exon usage during organoid formation. Number of alternative exon usage events detected using R-MATS in organoid and NBB compared to mESCs (red: exclusion, black: inclusion). (B) Pearson correlation of incLevelDifference obtained using R-MATS with the NBB and the organoids at days 7, 14 and 21.

**Figure S4.**
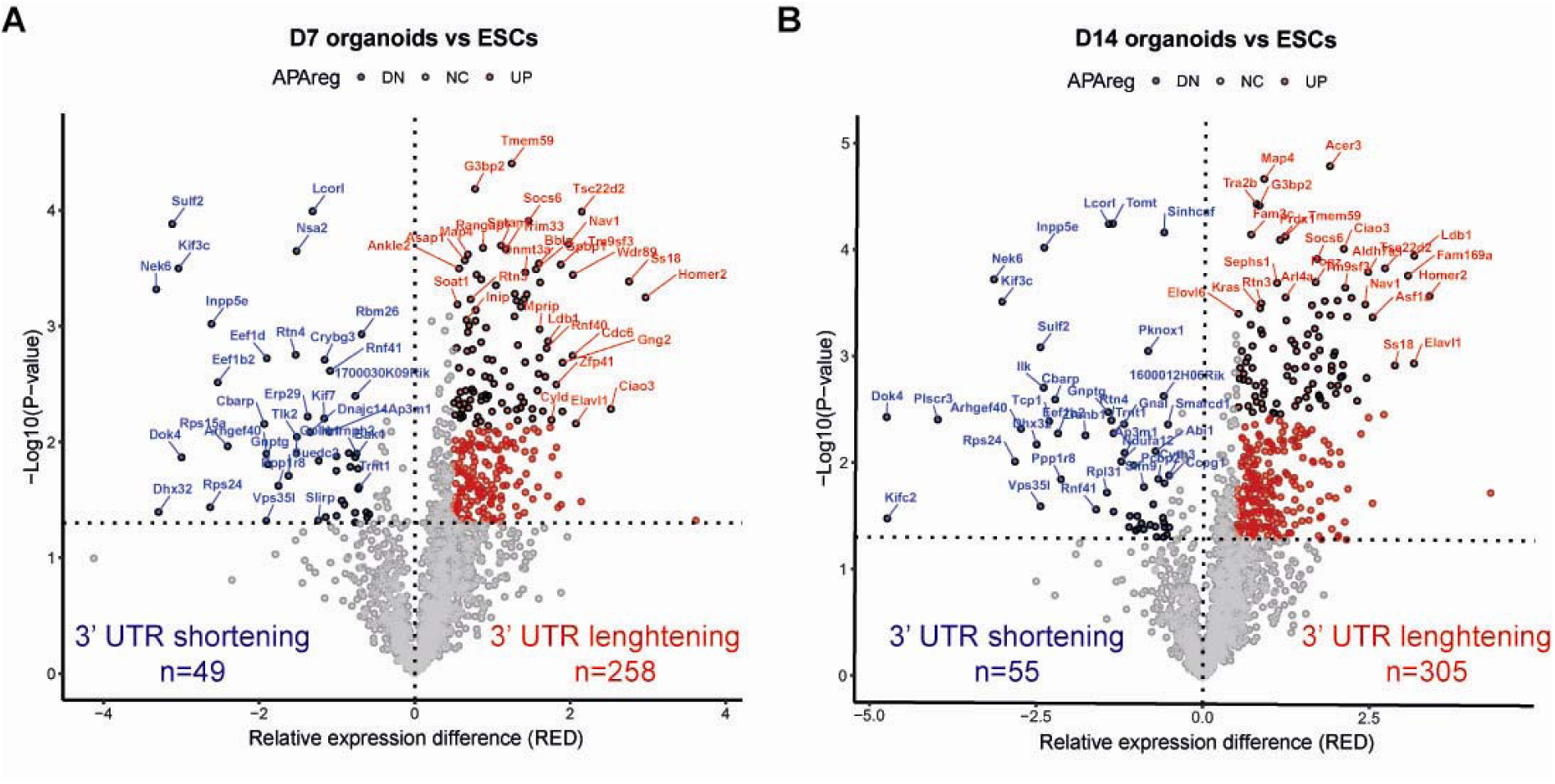
Evolution of alternative polyadenylation site usages during organoid formation. (A) Volcano plot of the distribution of genes identified as displaying proximal (blue) or distal (red) polyadenylation site usage in D7 organoids compared to mESCs. Genes were identified using APA analyser with the following cut-offs: -Log10(value>1.3) and RED (Relative Expression Difference) between proximal and distal polyadenylation site usage>0.5. (B) Same as in A, but for the D14 organoids compared to mESCs.

**Figure S5.**
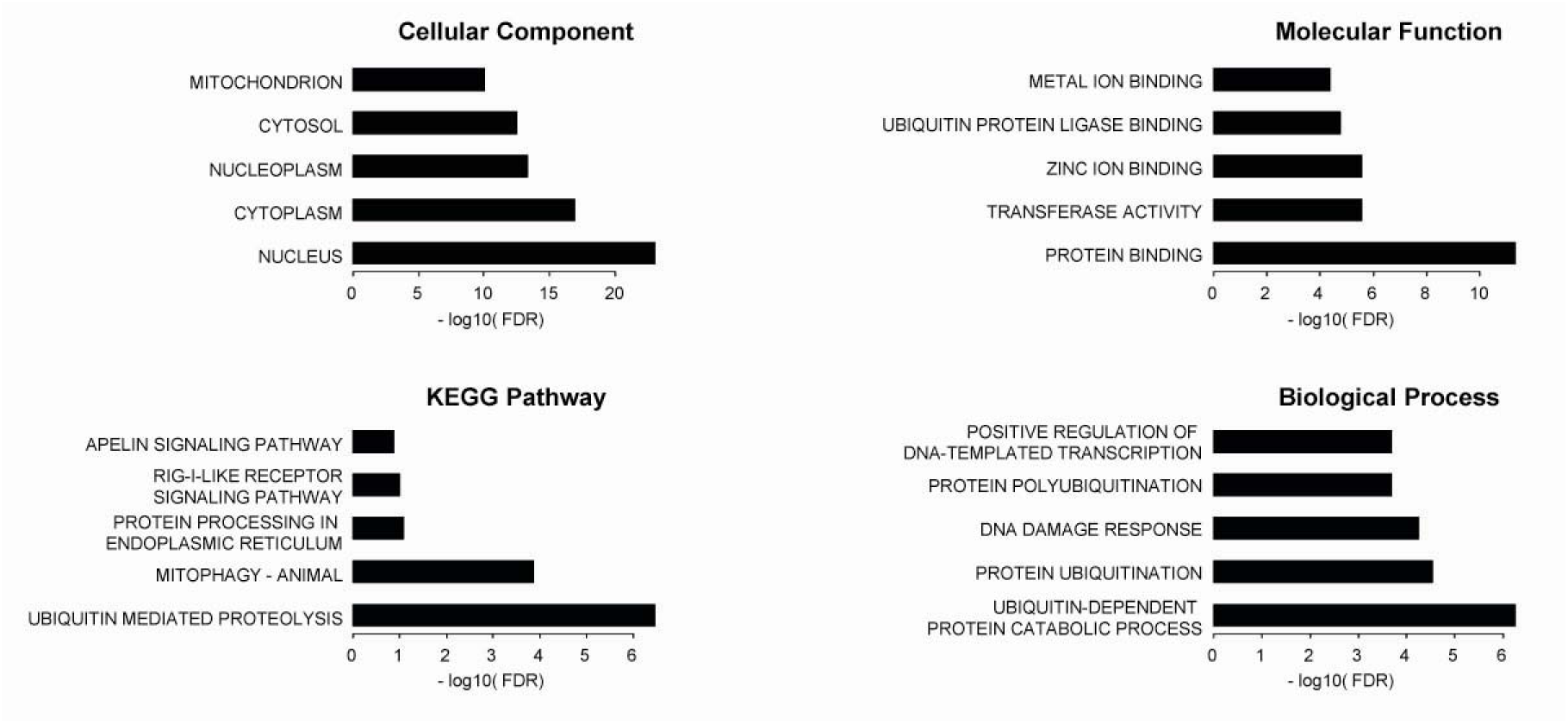
Gene ontology analysis on the genes with 3’UTR lengthening during organoid formation. Gene Ontology (GO) was performed using DAVID on the 366 genes with 3’UTR lengthening identified by APAnalyzer during the transition from mESCs to D21 brain organoids. For each GO term category, the five highest terms are shown for the differentially expressed gene lists. Black bars represent the –log10(FDR).

## Methods

### Differentiation of mouse embryonic stem cells into brain organoids

Mouse hybrid embryonic stem cells (mESCs) obtained from crosses between C57BL/6J and Mus musculus molossinus JF1 were expanded on Matrigel-coated dishes in ESGRO Complete Plus medium (Millipore) supplemented with a GSK3β inhibitor and penicillin/streptomycin as described [18]. For cortical organoid generation, ESCs were dissociated using Accumax and resuspended at a density of 30,000 cells/ml in Cortex Medium I: DMEM/F-12/GlutaMAX supplemented with 10% KSR, 0.1 mM of non-essential amino acids, 1 mM of sodium pyruvate, 0.1 mM of 2-mercaptoethanol, 100 U/mL penicillin/streptomycin, 1 µM DMH1-HCl and 240 nM IWP-2 [16]. Cells were seeded into 96-well ultra-low-adhesion plates (Sumitomo) at 3,000 ESCs per well in 100 µl. On day 4, an additional 100 µl of Cortex Medium I was added to each well. On day 7, 4 to 6 aggregates were transferred to 6-well low-adhesion plates and cultured in Cortex Medium 2: DMEM/F-12/GlutaMAX supplemented with N2 and B27 without vitamin A, 500 µg/mL of BSA, 0.1 mM of non-essential amino acids, 1 mM of sodium pyruvate, 0.1 mM of 2-mercaptoethanol, and 50 U/mL penicillin/streptomycin on an agitating platform. On day 14, 2 ml of medium was replaced with 2.5 mL of fresh Cortex Medium 2. Cultures were routinely tested for the absence of mycoplasma.

### Newborn brain samples

Newborn brains were collected from 1- to 3-day-old mice obtained from reciprocal crosses between C57BL/6J and Mus musculus molossinus JF1. Immediately after dissection, tissues were snap-frozen in liquid nitrogen and stored at −80 °C.

### Fixation, cryoprotection, cryosectioning and immunofluorescence on brain organoids

Organoids were processed as described by Madeline Lancaster’s lab (https://www2.mrc-lmb.cam.ac.uk/groups/lancaster/), with slight modifications. Briefly, organoids were fixed in 4% PFA for 15 min at RT and washed three times in PBS1X for 10 min. Samples were cryoprotected by incubation in 30% sucrose overnight at 4°C. Organoids were then incubated for 15 min at 37°C and embedded in 7.5% gelatin and 30% sucrose. Blocks were frozen on dry ice pellets and stored at −80°C until sectioning. For immunofluorescence experiments, cryosections (10µm) were air-dried for 5 minutes, rinsed in PBS, and blocked/permeabilised in PBS containing 0.25% Triton X-100 and 4% horse serum for one hour at RT. Primary antibodies were incubated overnight at 4°C. The primary antibodies were (species, provider; catalog number): anti-NESTIN –Rat-401- (mouse, Santa-Cruz, sc-33677); PAX6 (Rabbit, Covance, PRB-278P); NANOG (mouse, BD Pharmingen, 560259); POU5F1 (rabbit, Cell Signalling, 2840), ELAVL3 (mouse, Santa Cruz, sc-515624), TUBB3 (mouse, Covance, MMS-435P); TBR1 (rabbit, Cell Signalling, 49661), REELIN (mouse, Covance; MAB5364), GRM5 (rabbit, Millipore, AB5675), and GFAP (rabbit, Dako, Z0334). Cryosections were washed three times with PBS and incubated with secondary antibodies (anti-mouse or anti-rabbit coupled to Cy2 or Cy3 with minimal cross-species reactivity, Jackson Immunoresearch) for 2h at RT, both diluted in PBS containing 0.1% Triton X-100 and 4% FBS. Nuclei were stained with DAPI (1:200, 5 min at RT). The slides were mounted with a drop of mowiol and observed under a fluorescent microscope with an Apotome mode (AxioImager Z1, Zeiss) or a confocal microscope (LSM980 AIRYSCAN, Zeiss).

### Transcriptomics

RNA Samples (brain organoids at day 7: n=2, day 14: n=2, day 21: n=2 and newborn brain:n=2) were extracted using the MACHEREY-NAGEL NucleoSpin RNA Plus kit. After DNase treatment with the Qiagen RNase-Free DNase Set kit, the RNA samples were cleaned up with the Qiagen RNeasy Mini kit. Library preparation, with the NEBNext Ultra II mRNA-Seq Kit and sequencing on a NovaSeqX+ apparatus were performed by IntegraGen SA according to the manufacturer’s protocol. Stranded paired-end RNA-seq reads were mapped to the mouse genome mm39, which was masked for B6/JF1 SNPs, using Hisat2 and the list of known splice sites produced with the GENCODE vM37 basic annotation (version 2.2.1; option: -no-softclip -known-splicesite-infile). The RNA-seq alignments were filtered with samtools for mapping quality and reads mapped in proper pairs (version 1.21; view option: -f 2 -q 20). The strand-specific coverages of the RNA-seq data were generated using bamCoverage (version 3.5.6; option: -normalizeUsing RPKM -filterRNAstrand forward/reverse) and visualised on the UCSC genome browser. Strand-specific coverage of replicates was overlaid using the track collection builder tool of UCSC for genome exploration.

The RNA-seq alignments were processed using htseq-count(version 2.0.9) with the GENCODE vM37 basic annotation to compute the read counts per gene (option: -s reverse). Differential gene expression analyses between samples were conducted on read counts using the DESeq2 R packages, version 1.38.3 [58]. Genes were considered as differentially expressed between groups when |log2-fold change| >2 with an adjusted P-value <0.05.

The read counts data were normalised with the rlog function from the DESeq2 package before data visualisation with a heatmap (pheatmap R packages version1.0.13) and PCA (plotPCA function from the DESeq2 package).

To identify differential alternative splicing events between two sample groups, the RNA-seq alignments were analysed using the rMATS turbo (version 4.3.0) and the GENCODE vM37 basic annotation (options: -libType fr-firststrand -t paired -variable-read-length -readLength 100) [19]. Differential alternative splicing events were considered when | IncLevelDifference| >0.2 and FDR <0.05. Differential splicing event positions were loaded on UCSC in bed format in order to visualise the effect on RNA-seq coverages.

To identify alternative polyadenylation, RNA-seq alignments were treated with the Apalyzer R packages (version 1.12.0) [23]. 3 ‘UTR polyadenylation sites reference was built using the GENCODE vM37 basic annotation, and significantly alternative polyadenylation events in 3’UTRs were called with APAdiff function (option CUTreads=10) when |RED|> 0.5 and p-value <0.05. To evaluate locally or globally alternative polyadenylation on RNA-seq coverage, coordinates of alternative 3′UTR were loaded on UCSC or were treated with the Deeptools suite (version 3.5.6). Briefly, computeMatrix tools were used in scale-regions mode (-a 500 -b 500 -m 2000 --skipZeros) to calculate the strand-specific coverage of the RNA-seq per alternative 3′UTR. computeMatrixOperations was used to filter and merge the matrix produced with plus or minus strand coverage, according to the strand of the alternative 3’UTR. The corresponding heatmap was produced with the plotHeatmap tool.

The functional annotation tool of DAVID was used to identify enriched biological terms for gene lists [59].

Statistical analyses, plots, heatmap and Venn Diagram (venneuler version 1.1-4) were generated with R version 4.2.2. RNA-seq data for ESCs were obtained from the GEO database GSE244146.

### Proteomics

Samples were rinsed with PBS and kept at -80°C until analysis. Samples (ESCs: n=4, brain organoids at day7: n=3, brain organoids at day14: n=4, and brain organoids at day21: n=4) were digested with trypsin using the Strap (Supspension trapping) protocol. The resulting peptides were resuspended in buffer A (0.05% TFA, 2% acetonitrile) and injected for online analysis using nanoflow HPLC (RSLC U3000, Thermo Fisher Scientific) coupled to a mass spectrometer (Exploris 480). Peptides were separated on a capillary column (0.075 mm × 250 mm, Acclaim Pepmap 100, reverse phase C18, NanoViper, Thermo Fisher Scientific) following a gradient of 2-40% buffer B in 120 min (A = 0.1% formic acid, B = 0.1% formic acid, 80% acetonitrile) at a flow rate of 300 nl/min. Spectra were recorded using Xcalibur 4.2 software (Thermo Fisher Scientific) with the 120minDIA2.meth method. These instruments are checked regularly (cleaning, fluids, calibration). Spectral data were analysed with DIANN 1.8.2b27 and Perseus v1.6.15.0, using the DIAGui1.4.2 Rscript. We used as database(s): RefProt_Mouse_2025_03_FPPCON_v1.8.2.predicted.speclib with the modifications: Carbamidomethylation, N-Term M excision. The entire experimental process was verified by K.E.K and S.U. The Exploris 480 was last visited by the manufacturer in January 2025. Proteomic data were analysed using the Perseus computational tool [60] –version 1.6.15.0-. Gene ontology was performed using ShinyGO [26]. The heatmaps were generated using Morpheus, https://software.broadinstitute.org/morpheus.

## Acknowledgement

We thank the members of the Arnaud lab, Journot lab and the Proteomics facility for feedback and discussion. We thank Pedro Miura for helpful advice on alternative polyadenylation. We thank Anne Guillou for training and access to the cryostat facility (RHEM), the Light Microscopy facility of Biocampus for training and access to the confocal microscope, and Nicholas Boucharel (IGF) for performing mycoplasma assays. We thank Nathalie Bouquier and Julie Perroy for providing the GRM5 antibody. We thank members of the iGReD Bioinformatics Platform (BIM) for their technical assistance and discussions on RNA sequencing analysis. This work was supported by the Grant CORGI (ANR-23-CE12-0017, to P.A. and T.B.).

## Author Contributions

Conceptualisation, B.M., P.A., S.U., F.C., and T.B.; methodology, B.M., P.A., S.U., F.C., and T.B.; investigation, A-C. F., C. G. G., K.E.K., S. P., S.U., F.C, and T.B.; writing of the original draft, A-C. F., and T.B.; writing: review & editing, A-C. F., B.M., P.A., S.U., F.C, and T.B.; funding acquisition, P.A. and T.B.; resources, P.A., S.U., F.C, and T.B.; supervision, S.U., F.C, and T.B. All authors read and approved the final manuscript.

## Declarations

### Ethical Approval

Not applicable.

### Consent for Publication

Not applicable

## Supplementary material

